# An integrative and conjugative element encodes an abortive infection system to protect host cells from predation by a bacteriophage

**DOI:** 10.1101/2020.12.13.422588

**Authors:** Christopher M. Johnson, M. Michael Harden, Alan D. Grossman

## Abstract

Bacteriophages and integrative and conjugative elements (ICEs, a.k.a. conjugative transposons) are mobile genetic elements that can move from one bacterial cell to another and often coexist in a host genome. Many bacterial species contain at least one temperate (lysogenic) phage and an integrative and conjugative element. Most strains of *Bacillus subtilis* are lysogenic for the phage SPß and also contain the integrative and conjugative element ICE*Bs1*. We found that the presence of ICE*Bs1* in cells inhibited production of SPß, during both activation of a lysogen and following *de novo* infection. The ICE*Bs1* gene *yddK (*renamed *spbK* for SPß killing) was both necessary and sufficient for the anti-SPß activity. Co-expression of *spbK* and *yonE*, in the absence of other ICE*Bs1* and SPß genes, resulted in inhibition of cell growth and loss of cell viability. These results indicate that together *spbK* and *yonE* affected a host process needed for normal growth and that *spbK* constitutes an abortive infection system. We found that this anti-SPß phenotype protected populations of *B. subtilis* from predation by SPß, likely providing selective pressure for the maintenance of ICE*Bs1* in *B. subtilis* populations*. spbK* encodes a TIR (Toll-interleukin-1 receptor)-domain protein with similarity to both plant antiviral proteins and animal innate immune signaling proteins. We postulate that selective many uncharacterized cargo genes in ICEs may confer selective advantage to cells by protecting against other mobile elements.

## Introduction

Mobile genetic elements can move between host genomes or within a host’s genome. The genomes of many bacterial species contain multiple functional and defective mobile elements, including insertion sequences, transposons, lysogenic phages, genomic islands, and integrative and conjugative elements (ICEs; also called conjugative transposons). In some cases, these elements constitute a substantial portion of the host genome (Perna *et al.*, 2001; Paulsen *et al.*, 2003; Sebaihia *et al.*, 2006; Darmon and Leach, 2014). Multiple elements within a given host have the potential to interact with each other, and likely co-evolve.

ICEs are mobile genetic elements that reside integrated in a host chromosome and are replicated, segregated, and passed to daughter cells along with the host genome (Wozniak and Waldor, 2010; Johnson and Grossman, 2015; Delavat *et al.*, 2017). Under certain conditions, or stochastically, an ICE can excise from the chromosome and transfer itself to a recipient cell via the element-encoded conjugation machinery, typically a type IV secretion system.

ICEs frequently carry cargo genes that are not essential for their own lifecycle, but instead benefit the host. Most well-studied ICEs were discovered because of such phenotypes (Johnson and Grossman, 2015). For example, the ICE Tn*916* was discovered because it confers tetracycline resistance to host cells and can move between cells via conjugation (Franke and Clewell, 1981a; Franke and Clewell, 1981b). Likewise, many other ICEs were identified because they carry genes that confer specific phenotypes including: antibiotic resistance (Shoemaker *et al.*, 1980; Stuy, 1980; Roberts and Smith, 1980; Nugent, 1981; Mays *et al.*, 1982; Magot, 1983), pathogenesis (Carter *et al.*, 2010), symbiosis (Ramsay *et al.*, 2006), and metabolic functions (Rauch and De Vos, 1992; Hochhut *et al.*, 1997; Ravatn *et al.*, 1998; Nishi *et al.*, 2000).

Many ICEs have been identified by means other than the phenotype conferred by their cargo genes. In these cases, the functions of the cargo genes are largely unknown. We suspect that many of these cargo genes encode functions that are beneficial to the host under certain conditions, but that the appropriate conditions have not been identified.

Many strains of *Bacillus subtilis* contain at least two functional mobile genetic elements, the integrative and conjugative element ICE*Bs1* (Auchtung *et al.*, 2005; Auchtung *et al.*, 2016) and the lysogenic phage SPß (Warner *et al.*, 1977). *B. subtilis* strains also contain several defective mobile genetic elements (Kunst *et al.*, 1997; Barbe *et al.*, 2009; Smith *et al.*, 2014). We found that ICE*Bs1* contains a cargo gene that protects cells from predation by the lysogenic phage SPß.

ICE*Bs1* (Fig. 1) is found integrated in the *B. subtilis* genome in *trnS-leu2*, the gene for a leucine-tRNA. While integrated, most ICE*Bs1* genes are repressed (Auchtung *et al.*, 2005; Auchtung *et al.*, 2007). ICE*Bs1* is activated during the *recA-*dependent SOS response to DNA damage or in the presence of *B. subtilis* cells that do not have the element (Auchtung *et al.*, 2005). Under these conditions, ICE*Bs1* gene expression is derepressed, the element excises from the chromosome and can transfer to an available recipient via the element-encoded conjugation machinery. ICE*Bs1* was identified because of homology to other ICEs (Burrus *et al.*, 2002) and because it is regulated by cell-cell signaling (Auchtung *et al.*, 2005). At the time of its discovery, it was not known if ICE*Bs1* conferred a beneficial phenotype to its host.

**Fig. 1.**
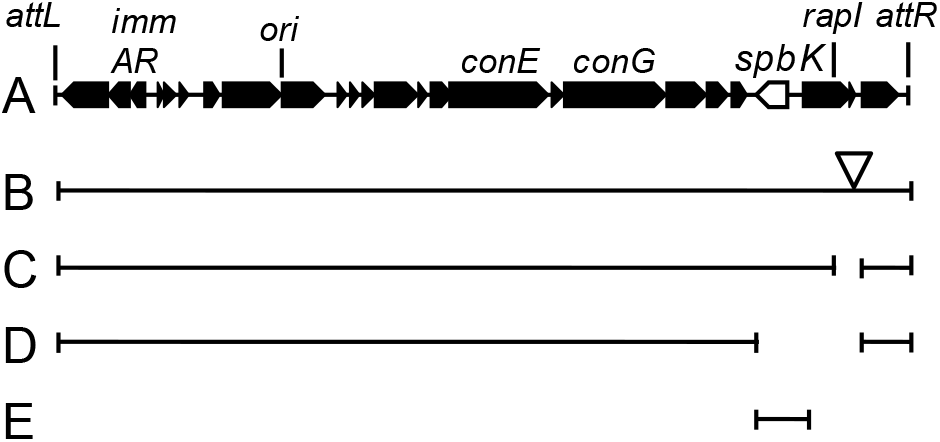
Map of ICE *Bs1* and some mutants. **A.** A linear map of ICE*Bs1* is shown. Genes are indicated as filled boxes with arrowheads at the ends indicating the direction of transcription. *spbK* is shown as an open arrow. The attachment sites *attL* and *attR* mark the junctions between ICE*Bs1* and chromosomal sequences. Below the map are shown the ICE*Bs1* mutants used in this work. The regions of ICE*Bs1* that are present in each construct are shown as bars beneath the map. **B.** ICE*Bs1*::*kan*. The open triangle indicates that this construct contains a *kan* gene inserted in the intergenic region between *rapI* and *yddM*. **C.** ICE*Bs1* Δ(*rapI-phrI*). **D.** ICE*Bs1* Δ(*spbK-phrI*). **E.** ICE*Bs1*^0^ *spbK*+. Many of these derivatives of ICE*Bs1* are used in multiple strains as indicated in Table 1.

**Table 1.**
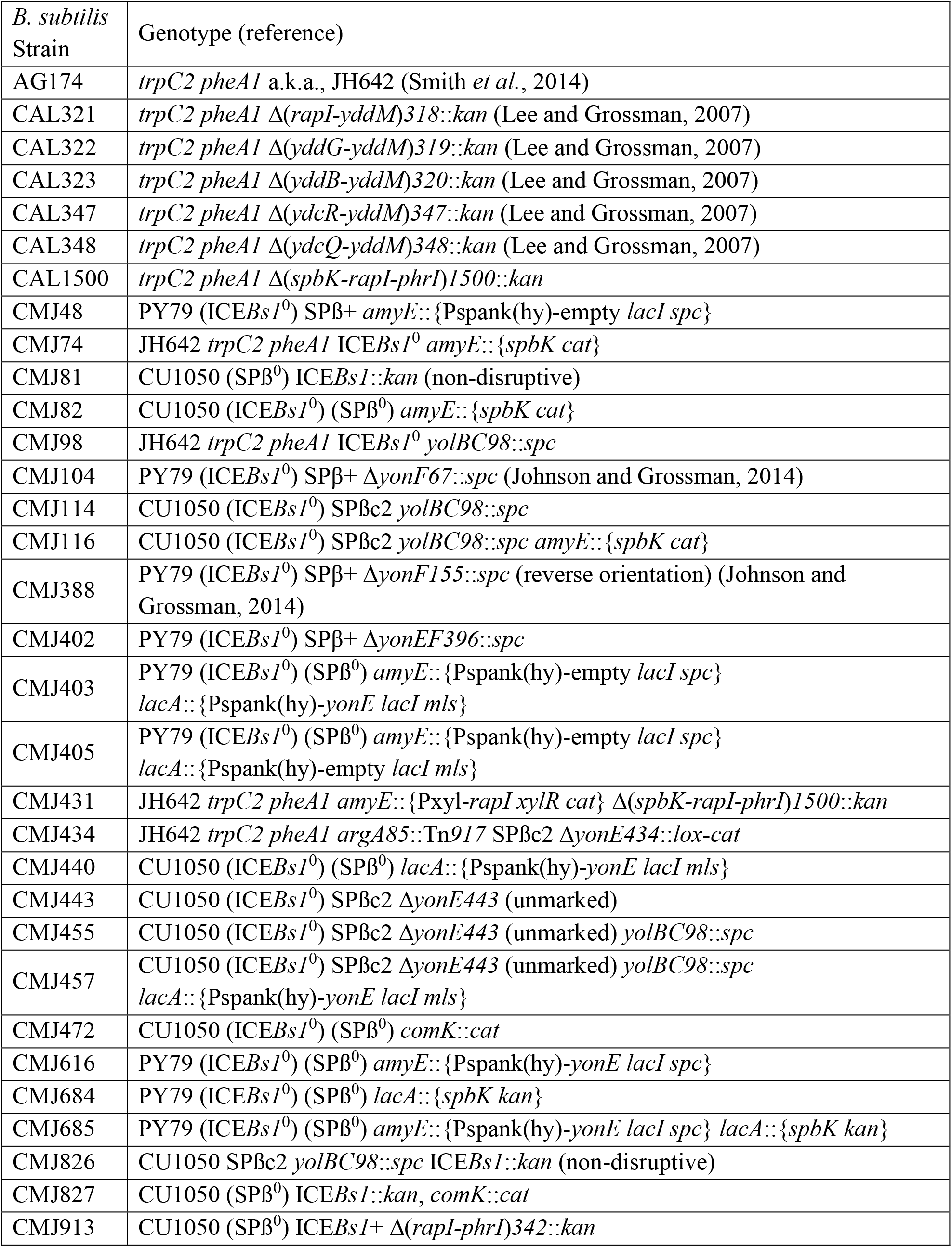
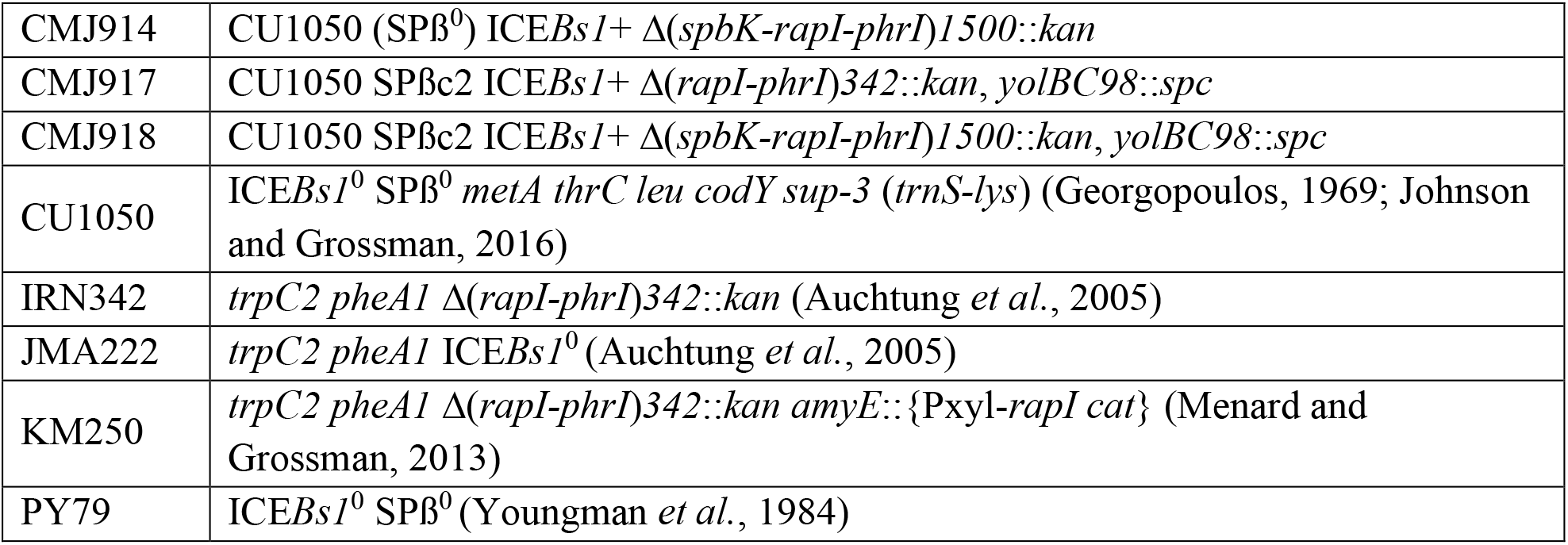
*B. subtilis* strains used.

| *B. subtilis* Strain | Genotype (reference) |
| --- | --- |
| AG174 | *trpC2 pheA1* a.k.a., JH642 (Smith *et al.*, 2014) |
| CAL321 | *trpC2 pheA1* $\Delta$(*rapI-yddM*)318::*kan* (Lee and Grossman, 2007) |
| CAL322 | *trpC2 pheA1* $\Delta$(*yddG-yddM*)319::*kan* (Lee and Grossman, 2007) |
| CAL323 | *trpC2 pheA1* $\Delta$(*yddB-yddM*)320::*kan* (Lee and Grossman, 2007) |
| CAL347 | *trpC2 pheA1* $\Delta$(*ydcR-yddM*)347::*kan* (Lee and Grossman, 2007) |
| CAL348 | *trpC2 pheA1* $\Delta$(*ydcQ-yddM*)348::*kan* (Lee and Grossman, 2007) |
| CAL1500 | *trpC2 pheA1* $\Delta$(*spbK-rapI-phrI*)1500::*kan* |
| CMJ48 | PY79 (ICEBsI<sup>0</sup>) SPβ+ *amyE*::{Pspank(hy)-empty *lacI spc*} |
| CMJ74 | JH642 *trpC2 pheA1* ICEBsI<sup>0</sup> *amyE*::{*spbK cat*} |
| CMJ81 | CU1050 (SPβ<sup>0</sup>) ICEBsI::*kan* (non-disruptive) |
| CMJ82 | CU1050 (ICEBsI<sup>0</sup>) (SPβ<sup>0</sup>) *amyE*::{*spbK cat*} |
| CMJ98 | JH642 *trpC2 pheA1* ICEBsI<sup>0</sup> *yolBC98*::*spc* |
| CMJ104 | PY79 (ICEBsI<sup>0</sup>) SPβ+ Δ*yonF67*::*spc* (Johnson and Grossman, 2014) |
| CMJ114 | CU1050 (ICEBsI<sup>0</sup>) SPβc2 *yolBC98*::*spc* |
| CMJ116 | CU1050 (ICEBsI<sup>0</sup>) SPβc2 *yolBC98*::*spc amyE*::{*spbK cat*} |
| CMJ388 | PY79 (ICEBsI<sup>0</sup>) SPβ+ Δ*yonF155*::*spc* (reverse orientation) (Johnson and Grossman, 2014) |
| CMJ402 | PY79 (ICEBsI<sup>0</sup>) SPβ+ Δ*yonEF396*::*spc* |
| CMJ403 | PY79 (ICEBsI<sup>0</sup>) (SPβ<sup>0</sup>) *amyE*::{Pspank(hy)-empty *lacI spc*}<br>*lacA*::{Pspank(hy)-*yonE lacI mls*} |
| CMJ405 | PY79 (ICEBsI<sup>0</sup>) (SPβ<sup>0</sup>) *amyE*::{Pspank(hy)-empty *lacI spc*}<br>*lacA*::{Pspank(hy)-empty *lacI mls*} |
| CMJ431 | JH642 *trpC2 pheA1 amyE*::{PxyI-*rapI xylR cat*} $\Delta$(*spbK-rapI-phrI*)1500::*kan* |
| CMJ434 | JH642 *trpC2 pheA1 argA85*::Tn917 SPβc2 Δ*yonE434*::*lox-cat* |
| CMJ440 | CU1050 (ICEBsI<sup>0</sup>) (SPβ<sup>0</sup>) *lacA*::{Pspank(hy)-*yonE lacI mls*} |
| CMJ443 | CU1050 (ICEBsI<sup>0</sup>) SPβc2 Δ*yonE443* (unmarked) |
| CMJ455 | CU1050 (ICEBsI<sup>0</sup>) SPβc2 Δ*yonE443* (unmarked) *yolBC98*::*spc* |
| CMJ457 | CU1050 (ICEBsI<sup>0</sup>) SPβc2 Δ*yonE443* (unmarked) *yolBC98*::*spc lacA*::{Pspank(hy)-*yonE lacI mls*} |
| CMJ472 | CU1050 (ICEBsI<sup>0</sup>) (SPβ<sup>0</sup>) *comK*::*cat* |
| CMJ616 | PY79 (ICEBsI<sup>0</sup>) (SPβ<sup>0</sup>) *amyE*::{Pspank(hy)-*yonE lacI spc*} |
| CMJ684 | PY79 (ICEBsI<sup>0</sup>) (SPβ<sup>0</sup>) *lacA*::{*spbK kan*} |
| CMJ685 | PY79 (ICEBsI<sup>0</sup>) (SPβ<sup>0</sup>) *amyE*::{Pspank(hy)-*yonE lacI spc*} *lacA*::{*spbK kan*} |
| CMJ826 | CU1050 SPβc2 *yolBC98*::*spc* ICEBsI::*kan* (non-disruptive) |
| CMJ827 | CU1050 (SPβ<sup>0</sup>) ICEBsI::*kan, comK*::*cat* |
| CMJ913 | CU1050 (SPβ<sup>0</sup>) ICEBsI+ $\Delta$(*rapI-phrI*)342::*kan* |
| CMJ914 | CU1050 (SPβ<sup>0</sup>) ICEBsI+ $\Delta$(*spbK-rapI-phrI*)1500::*kan* |
| CMJ917 | CU1050 SPβc2 ICEBsI+ $\Delta$(*rapI-phrI*)342::*kan*, *yolBC98*::*spc* |
| CMJ918 | CU1050 SPβc2 ICEBsI+ $\Delta$(*spbK-rapI-phrI*)1500::*kan*, *yolBC98*::*spc* |
| CU1050 | ICEBsI<sup>0</sup> SPβ<sup>0</sup> *metA thrC leu codY sup-3 (trnS-lys)* (Georgopoulos, 1969; Johnson and Grossman, 2016) |
| IRN342 | *trpC2 pheA1* $\Delta$(*rapI-phrI*)342::*kan* (Auchtung *et al.*, 2005) |
| JMA222 | *trpC2 pheA1* ICEBsI<sup>0</sup> (Auchtung *et al.*, 2005) |
| KM250 | *trpC2 pheA1* $\Delta$(*rapI-phrI*)342::*kan amyE*::{Pxyl-*rapI cat*} (Menard and Grossman, 2013) |
| PY79 | ICEBsI<sup>0</sup> SPβ<sup>0</sup> (Youngman *et al.*, 1984) |

SPß is a temperate phage of *B. subtilis*. In strains lysogenic for SPß the phage is integrated in *spsM*, near the terminus of replication (Zahler *et al.*, 1977; Lazarevic *et al.*, 1999; Abe *et al.*, 2017). SPß contains genes needed for production of and resistance to the peptide antibiotic sublancin (Paik *et al.*, 1998; Lazarevic *et al.*, 1999), providing a growth advantage to the host in the presence of cells sensitive to sublancin. Most phage genes are repressed in the lysogen, but during the *recA*-dependent SOS response to DNA damage, SPß gene expression is induced, the phage excises from the host chromosome, enters the lytic cycle producing progeny phage, and causes cell lysis and release of phage particles.

We found that the presence of ICE*Bs1* in *B. subtilis* inhibited production of active SPß, both when the phage was activated from the lysogenic state and during *de novo* infection. The ICE*Bs1* gene *spbK*, although dispensable for conjugation, was necessary and sufficient for the inhibition of SPß. The *spbK* gene product contains a Toll/Interleukin-1 Receptor (TIR) domain that was needed for function. The anti-SPß phenotype (abortive infection) caused by *spbK* was dependent on the SPß gene *yonE*. We found that *yonE* was essential for SPß lytic growth, but not for establishing a lysogen. Co-expression of *spbK* and *yonE* inhibited host cell growth and caused a drop in cell viability, even in the absence of any other ICE*Bs1* or SPß genes. The presence of ICE*Bs1* in cells prevented the spread of SPß, thereby protecting nearby *B. subtilis* cells from infection and allowing the population to continue growing. This phenotype likely provides strong selective pressure to maintain ICE*Bs1* in *B. subtilis*. We postulate that other ICEs might encode abortive infection, or other anti-phage systems, providing selective pressure for host cells to maintain these ICEs.

## Results

### ICE*Bs1* prevents SPß from forming plaques

ICE*Bs1* was not known to confer phenotypes to *B. subtilis*, aside from those directly related to conjugation. However, the left end of ICE*Bs1* (Fig. 1) encodes a phage-like repressor ImmR (Auchtung *et al.*, 2007), anti-repressor ImmA (Bose *et al.*, 2008), and recombinase Int (Lee *et al.*, 2007), leading us to wonder if ICE*Bs1* could provide immunity to phages that infect *B. subtilis*.

We tested if the presence of ICE*Bs1* in *B. subtilis* altered the ability of different phages to make plaques. We spread different phages on lawns of two *B. subtilis* indicator strains, one that was missing ICE*Bs1* and has been used as an indicator strain for SPß (CU1050; ICE*Bs1*^0^) (Zahler *et al.*, 1977; Johnson and Grossman, 2016) and an isogenic derivative that contained ICE*Bs1* (CMJ81; ICE*Bs1*+). SPß formed plaques on a lawn of the ICE*Bs1*^0^ strain (Fig. S1A), but not on a lawn of the isogenic ICE*Bs1+* strain (Fig. S1B), even when 100-fold more plaque forming units (PFUs) were mixed with cells (Fig. S1C). Based on these results, we conclude that something about ICE*Bs1* was inhibiting a step in plaque formation by SPß.

### ICE*Bs1* reduces phage production during infection

To quantify the effects of ICE*Bs1* on the production of SPβ, we measured the kinetics of phage production during a single round of infection (Fig. 2A). We mixed ~10^5^ PFUs of SPβ with ~10^7^ cells (MOI = 0.01) for 5 min at 37 °C, pelleted the cells by centrifugation, washed the cells to remove unattached phage, and resuspended the cells in LB medium at 37 °C to allow for phage growth. The initial number of infective cells in the medium was determined by measuring the number of infective centers (PFUs) following the initial adsorption, and new phage production was monitored by tracking the subsequent increase in infective centers. For a strain without ICE*Bs1*, the initial number of infective centers in the culture was about 90% of the initial number of phage used to inoculate the culture (Fig. 2A). The number of infective centers in the culture began to increase about 25 minutes after initial infection, and plateaued about 45 minutes after initial infection. This indicated that SPβ had an eclipse period of about 25 minutes (Fig. 2A). The burst size (number of phage produced per infective cell) was 20 ± 7, somewhat less than the previously reported burst size of about 30 phage (Warner *et al.*, 1977).

**Fig. 2.**
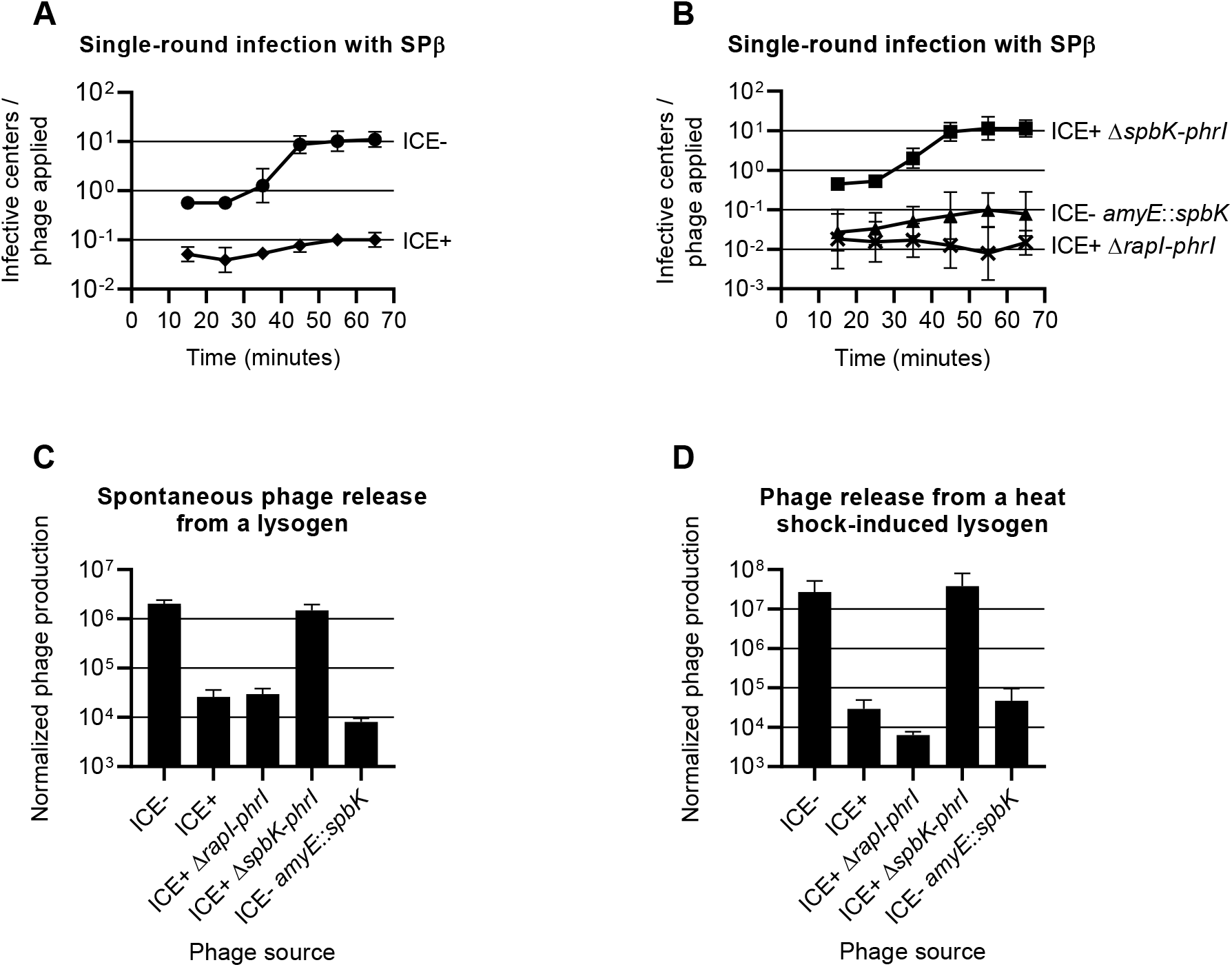
Production of SPß during the lytic cycle is reduced in cells containing ICE *Bs1* or *spbK*. **A.** Effect of ICE*Bs1* on single-round infection of *B. subtilis* cultures. SPß null strains were grown in rich medium, infected with phage (MOI = 0.01) and then diluted in fresh medium. The number of infective centers in the culture was tracked over time using strain CU1050 as the indicator (methods). circles, ICE*Bs1*^0^ (CU1050); diamonds, ICE*Bs1*^+^ (CMJ81). **B.** Effect of *spbK* on single-round infection of *B. subtilis* cultures. SPβ null strains were grown and infected with SPβ as in 2A. crosses, ICE*Bs1*^+^ Δ*rapI-phrI* (CMJ913); squares, ICE*Bs1*^+^ Δ(*spbK-rapI-phrI*) (CMJ914); triangles, ICE*Bs1*^0^ *amyE*::*spbK* ^(^CMJ82). **C.** Effect of ICE*Bs1* and *spbK* on spontaneous phage production. Strains carrying wild type SPß lysogens and different ICE*Bs1* variants were grown in rich medium; ICE*Bs1*^0^ (JMA222), ICE*Bs1*^+^ (AG174), ICE*Bs1*^+^ Δ*rapI-phrI* (IRN342), ICE*Bs1*^+^ Δ(*spbK-rapI-phrI*) (CAL1500), ICE*Bs1*^0^ *amyE::spbK* ^(^CMJ74). Supernatant was collected from each culture during exponential growth and used as a phage source in a plaque assay (methods). **D.** Effect of ICE*Bs1* and *spbK* on phage production after induction of a lysogen. Strains carrying ts SPß lysogens and different ICE*Bs1* variants were grown in rich medium; ICE*Bs1*^0^ (CMJ114), ICE*Bs1*^+^ (CMJ826), ICE*Bs1*^+^ Δ*rapI-phrI* (CMJ917), ICE*Bs1*^+^ *Δ(spbK-rapI-phrI*) (CMJ918), ICE*Bs1*^0^ *amyE*::*spbK* ^(^CMJ116). SPß lysogens were induced by a heat shock during exponential growth and supernatants were collected and used as a phage source in a plaque assay (methods). For C and D the Y axis shows the number of PFU/ml of culture divided by the OD600 of the culture.

Cells with ICE*Bs1* that were exposed to SPβ were less likely to become infective centers, and produced fewer phage per initial infective center. At an MOI of 0.01, the number of cells that produced any phage was reduced at least 10-fold relative to cells without ICE*Bs1* (Fig. 2A). Furthermore, the number of phage produced per initial infective center was ~2.2 ± 0.4 (Fig. 2A). Based on these results, we conclude that the presence of ICE*Bs1* in cells reduced the total number of phage released from the infected culture by at least 100-fold, or to about 0.1 progeny phage per infecting phage. This reduction will not support propagation of phage in the lytic cycle.

### ICE*Bs1* has little or no effect on entry of phage into cells

The ICE*Bs1*-dependent reduction in plaque formation and phage production could be due to reduced entry of phage into cells. Alternatively, a step in the phage lifecycle after entry could be inhibited. If the presence of ICE*Bs1* was causing a block in phage entry, then there should be a corresponding reduction in the frequency of lysogen formation. We used SPß that contained *spc*, conferring resistance to spectinomycin, to measure the frequency of lysogenization. Cells with or without ICE*Bs1* were mixed with SPß::*spc98* (MOI = 0.001), unbound phage were washed off, and cells were spread on plates containing spectinomycin to select for lysogens. The lysogenization frequency of cells without ICE*Bs1 (*CMJ472) was ~1% (1.1×10^−2^ ± 0.46×10^−2^), or approximately one lysogen per 100 initial phage. Similarly, the lysogenization frequency of ICE*Bs1*+ cells (CMJ827) was ~0.4% (4.3×10^−3^±1.3×10^−3^), or about 40% of that of the ICE*Bs1*^0^ cells. These results indicate that ICE*Bs1* has a relatively minor (if any) effect on lysogenization frequency and that the anti-SPß phenotype conferred by ICE*Bs1* was not due to a block in adhesion or entry of the phage.

### ICE*Bs1* reduces the number of phage released by SPß lysogens

We found that the presence of ICE*Bs1* in an SPß lysogen inhibited phage production. We grew lysogens in liquid medium and measured the number of PFUs present in the supernatant. We found that cultures of a lysogen without ICE*Bs1* had approximately 100-fold more PFUs/ml than cultures of an ICE+ lysogen (Fig. 2C). Together our results demonstrate that ICE*Bs1* acts primarily by blocking production of phage by infected cells, rather than by preventing infection of cells in the first place.

We also found that the presence of ICE*Bs1* prevented production of SPß following induction of a temperature sensitive lysogen. We grew strains with a temperature sensitive SPß lysogen (SPß*c2*) in rich medium, induced the lysogen by heat shock, and measured phage release. Phage production was reduced by ~1,000-fold in cells with ICE*Bs1* compared to cells without (Fig. 2D). Although production of functional phage particles was reduced, the cells were still killed following phage induction. Cell viability, as measured by colony forming units (CFUs), was reduced to ~0.1% after phage induction compared to right before phage induction for strains with (CMJ826) and without ICE*Bs1* (CMJ114). Based on these results we conclude that ICE*Bs1* was probably not preventing induction of SPß but rather was inhibiting production of active phage particles post-induction.

### The ICE*Bs1* gene *spbK* is necessary and sufficient to inhibit SPß

We were interested in determining which ICE*Bs1* gene was responsible for the inhibition of SPß. Most ICE*Bs1* genes are repressed when ICE*Bs1* is integrated in the host genome. Because the inhibition of SPß did not appear to depend on activation of ICE*Bs1*, we focused on the handful of ICE*Bs1* genes that are constitutively expressed, including genes toward the left and right ends of the element (Fig. 1). In preliminary testing of SPβ lysogens containing deletions of different regions of ICE*Bs1*, we found that strains in which *spbK* (formerly *yddK*) had been deleted did not inhibit phage production, whereas all strains in which *spbK* was intact inhibited production. Based on these results, we inferred that *spbK* was likely needed for ICE*Bs1*-mediated inhibition of spontaneous release of SPß from a lysogen.

We used three different assays to test the effects of *spbK* on SPβ. In all three assays, we compared three *B. subtilis* strains: an ICE*Bs1*+ strain with *spbK* (*Δ(rapI-phrI*)::*kan*, Fig. 1C), an ICE*Bs1*+ strain lacking *spbK* (*Δ(spbK-phrI*)::*kan*, Fig. 1D), and an ICE*Bs1*^0^ strain expressing *spbK* from its own promoter at an ectopic locus (ICE*Bs1*^0^ *amyE*::{*spbK kan*}, Fig. 1E). We measured: 1) the appearance of infective centers following a single round of infection with SPβ (Fig. 2B); 2) the number of phage spontaneously released from an SPβ lysogen (Fig. 2C); and 3) the number of phage produced after induction of a temperature sensitive SPβ lysogen (Fig. 2D). In all cases, we found that *spbK* was necessary for ICE*Bs1* to inhibit the formation of infective centers and the production of phage, and that ICE*Bs1*+ *ΔspbK* strains were indistinguishable from strains entirely lacking ICE*Bs1*. Furthermore, ectopic expression of *spbK* was sufficient to inhibit phage production in the absence of ICE*Bs1* in all three assays.

### Expression of the SPβ gene *yonE* inhibits acquisition of ICE*Bs1* and this inhibition is dependent on the ICE*Bs1* gene *spbK*

Based on the results described above, we thought that there might be at least one gene in SPß that was needed for the *spbK*-mediated inhibition of phage production. Results described below indicate that *yonE* is this SPß gene.

In previous work, we identified transposon insertions in recipients that reduced the ability of cells to acquire ICE*Bs1* from donor cells (Johnson and Grossman, 2014). Recipients were SPß lysogens, and we found that certain mutations in the SPβ gene *yonF* reduced the ability of would-be recipients to acquire ICE*Bs1* from donors in conjugation (Johnson and Grossman, 2014). Because the pattern of *yonF* mutations was not uniform, we hypothesized that the reduction in ICE*Bs1* acquisition was not due to loss of *yonF*, but rather due to increased expression *yonE*, the gene immediately downstream of *yonF*, driven by the promoter the drives transcription of the antibiotic resistance gene in the transposon.

We found that expression of *yonE* reduced the ability of cells to acquire a copy of ICE*Bs1*. We made a series of mutations in SPβ (Fig. 3A) and tested these for effects on the ability of cells to act as ICE*Bs1* recipients during conjugation. We found that an insertion of *spc* into a deletion of *yonF* (Δ*yonF*::*spc*) reduced acquisition of ICE*Bs1* only when *spc* was co-directional with *yonE*. Furthermore, deletion of *yonE* in this context eliminated the defect in acquisition of ICE*Bs1* (Fig. 3B). In the absence of all other SPß genes, expression of *yonE* from the IPTG-inducible promoter Pspank(hy) was sufficient to inhibit acquisition of ICE*Bs1* (Fig. 3B). We conclude that *yonE* in SPß is both necessary and sufficient to cause the decrease in acquisition of ICE*Bs1*.

**Fig. 3.**
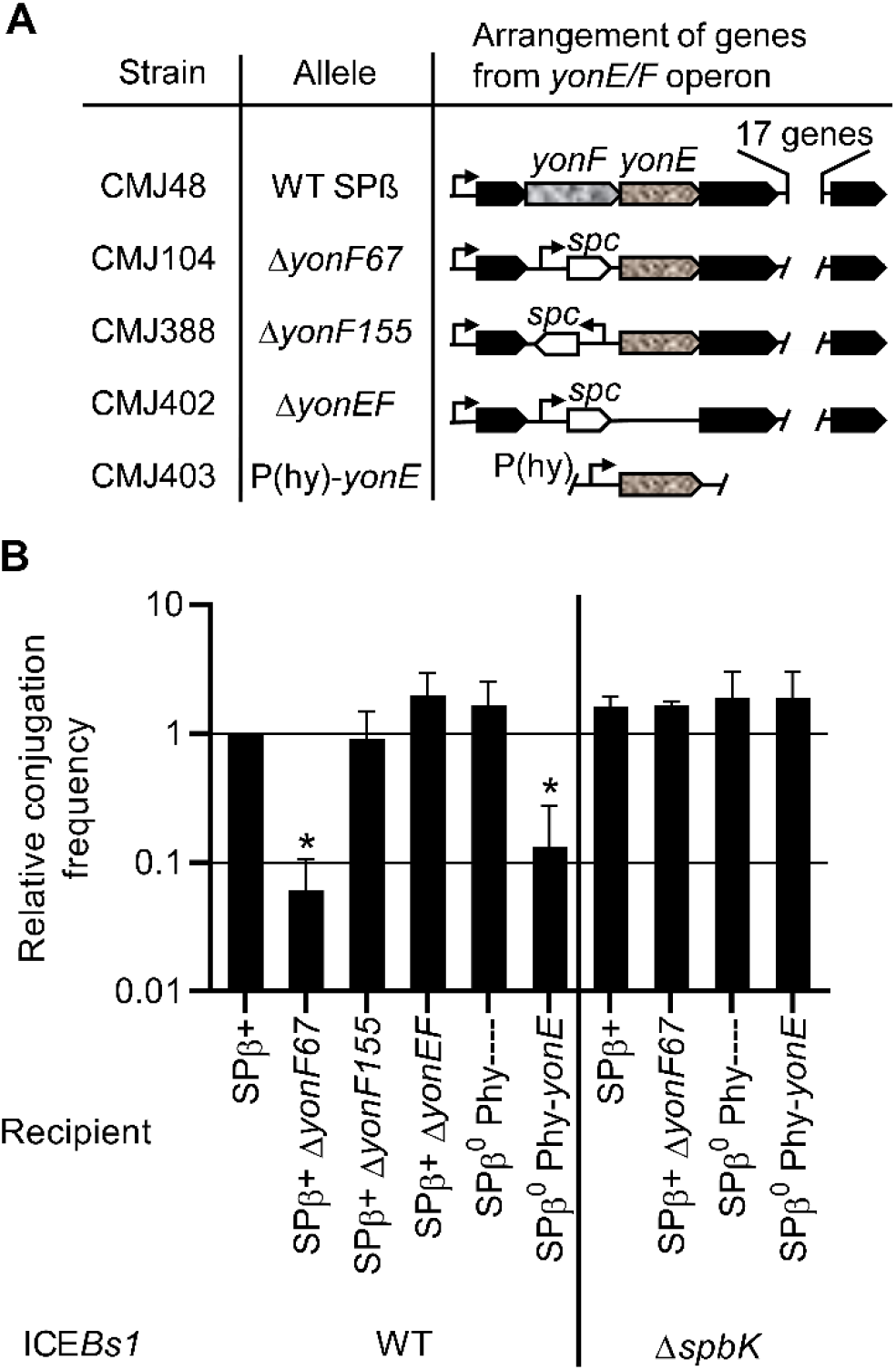
Expression of *yonE* in recipients reduces acquisition of ICE*Bs1* via conjugation. **A.** Map of the operon in SPß that contains *yonF* and *yonE*, and relevant mutations. Genes are shown as arrows. *yonF* and *yonE* are indicated by arrows filled with a mottled pattern. *spc* is shown as an open arrow. Promoters are shown as bent arrows. The allele and the recipient strain carrying that allele are indicated. **B.** The relative conjugation frequencies are shown, normalized to the conjugation frequency between a donor carrying a wild type ICE*Bs1* (KM250) and a recipient with a wild type SPß (CMJ48) within the same experiment, (average 9.7 x 10^−4^ ± 1.3 x 10^−3^ transconjugants/donor). The Δ*spbK* donor (CMJ431) carries an ICE*Bs1* in which *spbK-rapI-phrI* have been deleted. The recipient with the Pspank(hy) promoter and no *yonE* allele is CMJ405. Other recipient strain numbers are indicated in panel A. Each experiment was repeated ≥ 3 times. Asterisks indicate that the conjugation frequency with the given recipient is significantly different than that with the wild type control (p<0.05, t-test).

The decreased acquisition of ICE*Bs1* by recipients expressing *yonE* was dependent on the ICE*Bs1* gene *spbK*. We tested strains expressing *yonE* for the ability to acquire ICE*Bs1* that was missing *spbK* (ICE*Bs1* Δ*spbK*), and found that they all acquired the Δ*spbK* element at the same frequency as wild type recipients not expressing *yonE (*Fig. 3B, right end of panel). From these results we conclude that expression of *yonE* caused a defect in ICE*Bs1* acquisition and that this defect was dependent on the presence of *spbK* in the incoming ICE*Bs1.* We note that loss of *spbK* caused no reduction in conjugation efficiency (Fig. 3B), demonstrating that it is dispensable for conjugation.

### Co-expression of *yonE* and *spbK* causes a defect in cell growth and a drop in cell viability

We found that expression of *spbK (*from its own promoter) and *yonE (*from Pspank(hy)) together caused a severe growth defect. We grew cells containing both *spbK* and *yonE* in defined minimal medium and added IPTG (time = 0) to increase expression of *yonE* (Fig. 4A). This caused a rapid growth arrest as measured by optical density (Fig. 4A). In contrast, expression of either gene alone, *spbK* from its own promoter (*lacA*::*spbK*), or *yonE* from an inducible promoter (*amyE*::Pspank(hy)-*yonE*), had no obvious effect on growth (Fig. 4A). This growth arrest was accompanied by a ~1000-fold drop in viability as measured by plating for CFUs on LB plates made with Noble agar (Fig. 4B; see below). Together, these results indicate that co-expression of *yonE* and *spbK* is detrimental to cell growth.

**Fig. 4.**
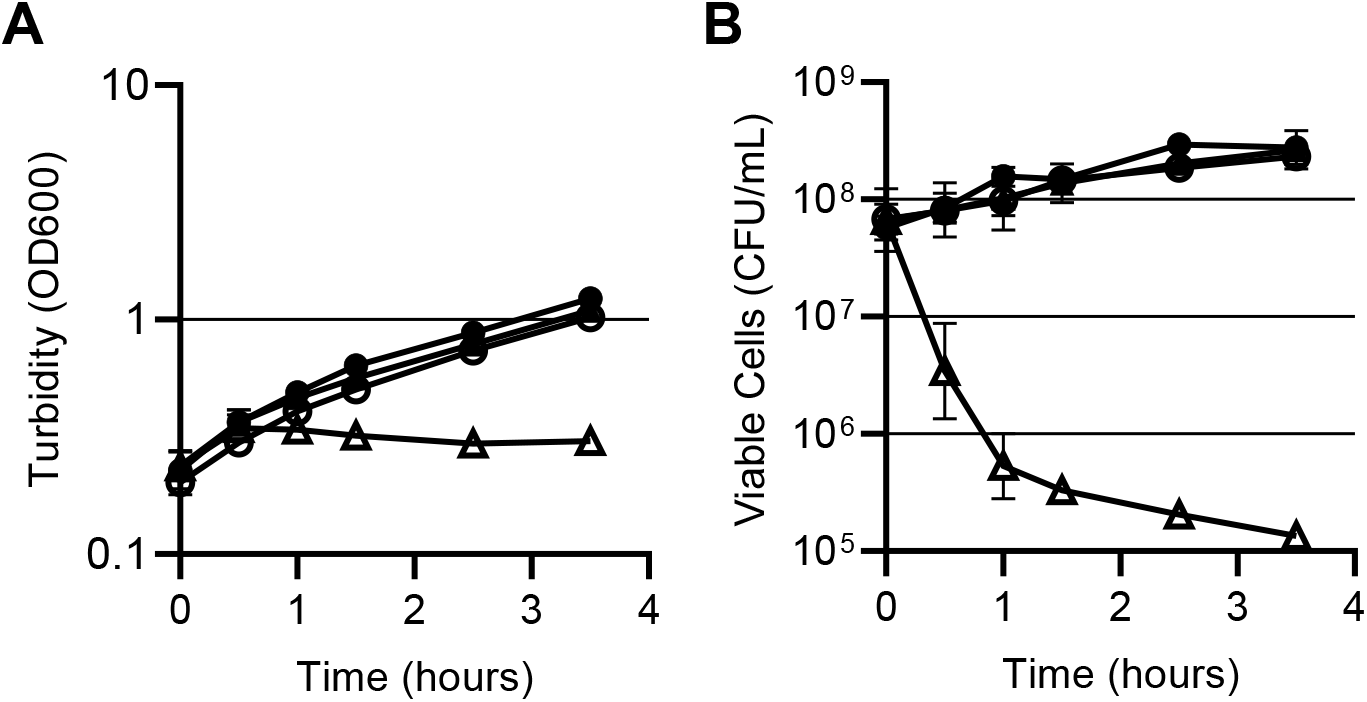
Co-expression of *spbK* and *yonE* kills cells and results in a growth defect. Strains null for ICE*Bs1* and SPß (closed circles, PY79), expressing *yonE* (closed triangles, *amyE*::Phy-*yonE*, CMJ616), expressing *spbK* (open circles, *lacA*::*spbK*, CMJ684), or both *yonE* and *spbK* (open triangles, CMJ685) were grown in minimal medium. *yonE* expression was induced by the addition of 1 mM IPTG and the culture turbidity (**A**), and cell viability (**B**) were followed over time. T = 0 samples were collected immediately prior to induction with IPTG. Cell viability was measured as the number of colony forming units per ml, measured by plating for CFUs on LB plates made with Noble agar. Experiment was repeated 3 times.

Despite growing normally in defined liquid medium prior to adding IPTG, cells containing both *lacA*::*spbK* and *amyE*::Pspank(hy)-*yonE* had a substantial plating defect (~200-fold) when plated on LB plates made with standard bacto-agar (Difco), and had a distinct small colony morphology even in the absence of IPTG. The plating and colony size defects were eliminated when the cells were plated on LB plates made with Noble agar (Difco) (Fig. S2), a purified form of agar that is used to culture some fastidious organisms. We hypothesize that a component of bacto-agar sensitizes cells to the detrimental impact of co-expressing *spbK* and *yonE*, such that leaky expression from Pspank(hy)-*yonE* is sufficient to trigger the growth defect.

### *yonE* is needed for phage production

To determine the effect of *yonE* on phage production, we made an unmarked deletion of *yonE* (Δ*yonE443*) in a temperature-sensitive SPß lysogen. We found that cultures of this inducible Δ*yonE* lysogen cleared comparably to a *yonE*+ strain following a shift to high temperature (Fig. 5A), demonstrating that *yonE* is not needed for induction of SPβ from a lysogenic state, nor is it needed to cause host cell lysis.

**Fig 5.**
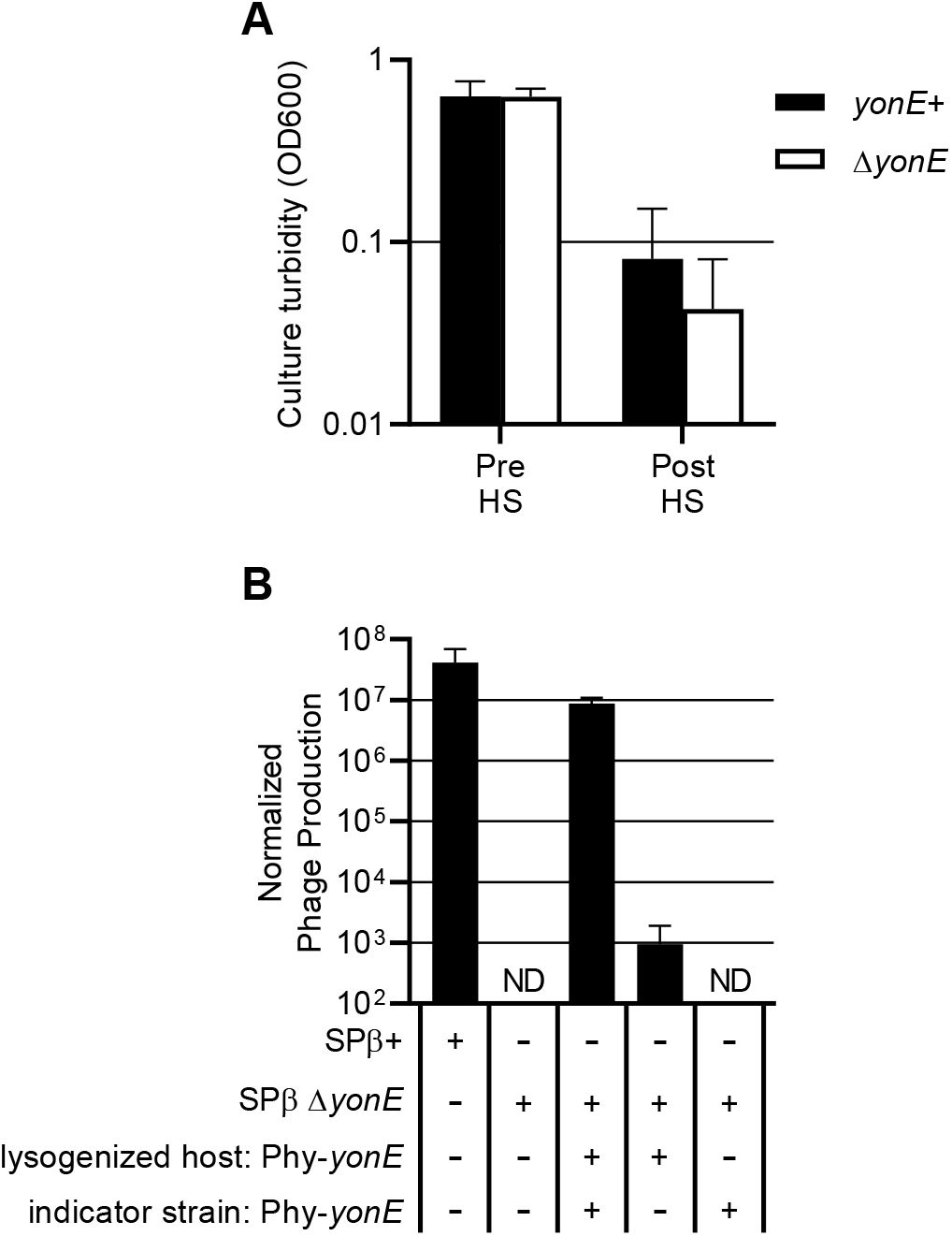
*yonE* is needed for production of infectious phage. **A.** Strains carrying a temperature-sensitive SPß lysogen with a wild type *yonE* allele (black bars, CMJ114) or Δ*yonE* (white bars, CMJ455) were cultured in rich medium, then heat shocked for 20 minutes (methods). The Y axis shows the average and standard deviation of the OD600 from each culture immediately before and 70 minutes (± 5 minutes) after the heat shock. Each experiment was repeated ≥3 times. **B.** Phage were prepared by culturing strains with a temperature sensitive SPß lysogen with a wild type *yonE* allele (CMJ114), Δ*yonE* (CMJ455), or a Δ*yonE* allele with *yonE* complemented from the chromosome (*lacA::*Phy-*yonE*, CMJ457) to an OD600 of approximately 0.4, and then heat shocking the cultures and collecting phage (methods). Lysates were then spread on lawns of the indicator strain CU1050 or an indicator with *lacA::*Phy-*yonE* (CMJ440) and incubated overnight to allow plaque formation. The Y-axis shows the average and standard deviation of the number of infectious phage /ml, normalized by dividing by the OD600 of the culture at the time of heat shock. ND = not detected. Each experiment was repeated ≥3 times.

Despite the fact that the Δ*yonE* host cells lysed, there were no detectable viable phage (< 10 PFUs/ml) produced by the mutant lysogen (Fig. 5B, first two columns). This defect in phage production was partially complemented by expression of *yonE* from an ectopic chromosomal locus. These Δ*yonE* phage (recovered from the complemented strain) were capable of forming plaques on an indicator strain that also expressed *yonE* (Fig. 5B, last three columns). Although a small number of phage produced by a Δ*yonE* lysogen were able to form plaques on a *yonE*-indicator strain, analysis of lysogenized cells obtained from these plaques revealed that the phage had a wild type copy of *yonE*, likely obtained through homologous recombination with the *yonE* allele on the chromosome of the original host strain. Based on these results, we conclude that *yonE* is essential for production of SPβ.

To determine if *yonE* is needed to form a lysogen, we made a stock of *spc*-marked Δ*yonE* phage by growing the Δ*yonE* mutant on a *B. subtilis* strain ectopically expressing *yonE*. The frequency of lysogenization of *spc*-marked *yonE*+ and Δ*yonE* phage were both approximately 1%, indicating that *yonE* is not needed for lysogen formation.

The function of *yonE* is not known. However, there are homologs in other phages, including the phage C-ST from *Clostridium botulinum*. The region of homology between C-ST and SPß extends from *yonG* to *yomZ*, indicating that these genes may encode conserved phage functions (Sakaguchi *et al.*, 2005). The phage E3 from *Geobacillus* encodes a putative portal protein (accession number AJA41333) that is 25% identical to YonE (van Zyl *et al.*, 2015). Portal proteins are one of three molecular components involved in packaging the phage genome into the capsid during maturation. The other two components are the large and small terminase subunits (Rao and Feiss, 2015). Additional homology searches using NCBI BLAST revealed that YonF is a member of the terminase 1 superfamily and encodes a terminase 6 multidomain, typical of large terminase subunit proteins. These results indicate that YonE and YonF may be a part of the SPß head packaging machinery. This notion is consistent with the need for *yonE* in production of functional phage, but not in host cell lysis or formation of lysogens.

### SpbK contains a TIR domain involved in protein-protein interaction

*spbK* is predicted to encode a 266 amino acid protein. Using the NCBI Delta-BLAST search tool (Boratyn *et al.*, 2012) we found that the C-terminal region of SpbK (amino acids 113-266) contains a Toll Interleukin-1 Receptor (TIR) domain in the TIR_2 superfamily (accession: cl23749) (Fig. S3A). Proteins containing TIR domains have been found in animals, plants, and bacteria. In animals, such proteins are involved in signaling cascades in development and in immune activation (Narayanan and Park, 2015). In plants they mediate disease resistance, often in response to infectious agents (McHale *et al.*, 2006). Some pathogenic bacteria encode TIR domain proteins that interact with eukaryotic host TIR domain proteins to modulate the host immune response (Rana *et al.*, 2013). Many non-pathogenic bacteria also contain TIR domain proteins and it is thought that the TIR domains mediate protein-protein interaction (Spear *et al.*, 2009). Recent work has also implicated some bacterial TIR domain proteins as components of anti-phage defense systems, though the mechanism of defense is not understood (Doron *et al.*, 2018).

Where they have been studied, TIR domains mediate protein-protein interactions by interacting with other TIR domains. *spbK* is the only gene in *B. subtilis*, including all horizontally acquired sequences (e.g, ICE*Bs1* and SPß), predicted to encode a TIR domain. Using a yeast two-hybrid assay, we found that full-length SpbK multimerizes *in vivo* (Fig. S2B). Additionally, we found that the TIR domain alone interacted with both full-length SpbK and with the TIR domain, but that deleting the TIR domain abolished all interaction between SbpK proteins (Fig. S2B). We also tested for, but were unable to detect, interaction between SpbK and YonE.

### ICE*Bs1* protects *B. subtilis* populations from attack by SPß

Cells containing ICE*Bs1* and SPβ die when SPß enters the lytic cycle, and very few infectious phage are produced. We found that in a population of cells, this abortive infection system in ICE*Bs1* protected cells from killing by SPß. We grew SPß-cured strains of *B. subtilis* that either contained or did not contain ICE*Bs1*, infected the cultures with SPß at a low multiplicity of infection (MOI = 0.01), and tracked the growth (optical density) of the culture, the concentration of viable cells (including lysogens), and free phage over time (Fig. 6). When cultures of an ICE*Bs1*^0^ strain were infected with a clear plaque mutant of SPß (incapable of forming lysogens) the cells continued to grow at the same rate as an uninfected culture for approximately 1.5 hours, then the majority of the cells abruptly died, as evidenced by a decrease in optical density (Fig. 6A) and an approximately 5,000-fold decrease in CFUs (Fig. 6B). During this time (1.5 hrs) the concentration of phage in the culture (Fig. 6C) surpassed the concentration of cells (Fig. 6B).

**Fig. 6.**
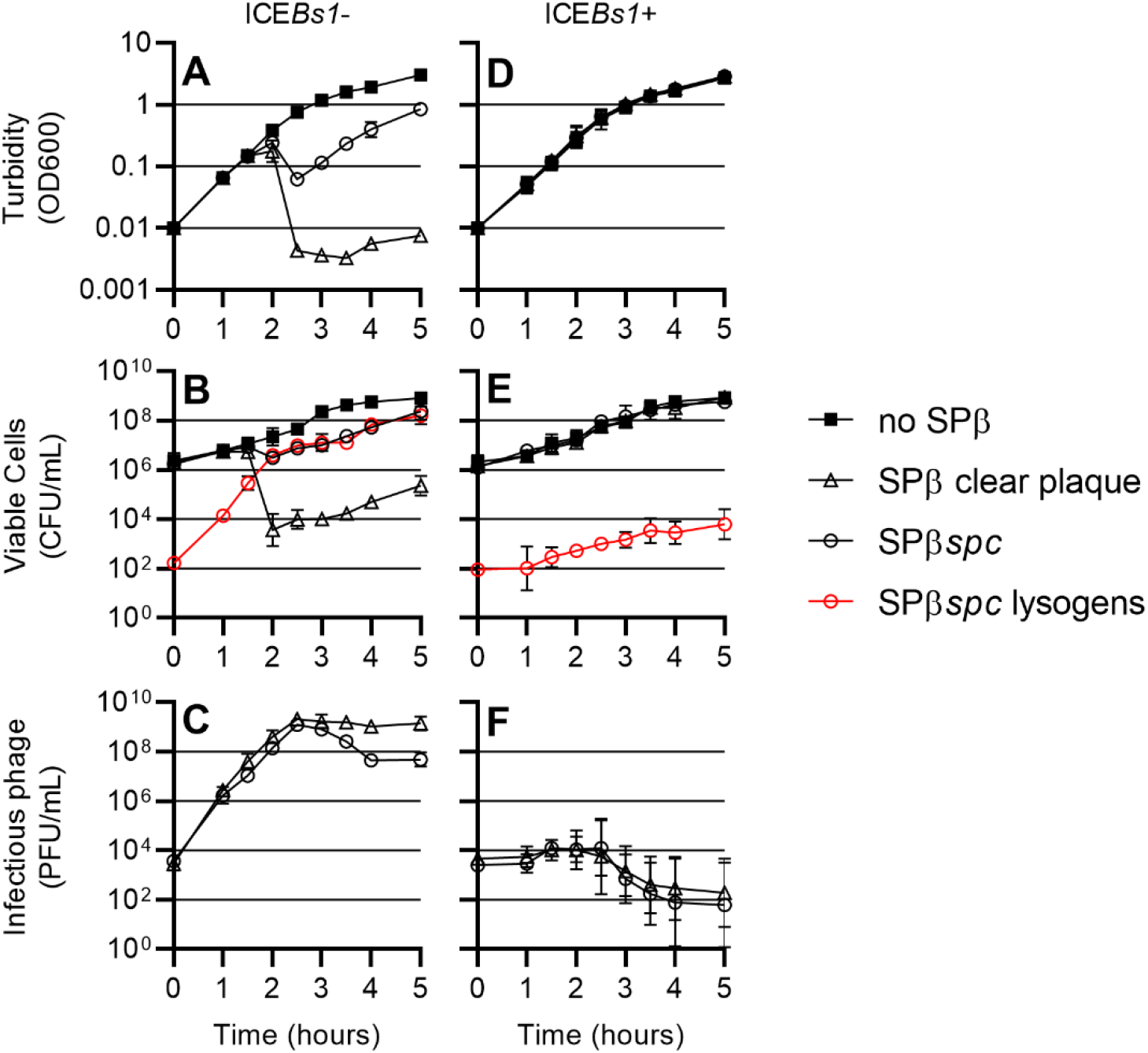
ICE*Bs1* protects *B. subtilis* populations against SPß. **A, B, C,** SPß^0^ ICE*Bs1*^0^ (CU1050) and **D, E, F,** SPß^0^ ICE*Bs1+ (*CMJ81) were grown in rich medium, infected with no phage (filled squares), SPß clear plaque (open triangles), or SPß::*spc* (open circles) at an MOI of approximately 1:100 and diluted in fresh medium to an OD600 of Culture turbidity (**A, D**), CFUs/ml (**B, E)**, and the number of infectious phage per ml of culture (**C, D**) were tracked over time. Additionally, for cultures infected with SPß::*spc*, the number of lysogenized cells were tracked over time (red open circles).

In contrast, when an ICE*Bs1*+ strain was infected with the clear plaque mutant of SPß (MOI = 0.01), cell growth was indistinguishable from an uninfected culture as measured by both the optical density (Fig. 6D) and the number of CFUs (Fig. 6E). The population of phage in the culture generally decreased to below the initial inoculum (Fig. 6F). These results indicate that the presence of ICE*Bs1* is beneficial to the population of cells even though individual infected cells may not survive.

Experiments described above were done with SPß phage that were unable to form lysogens. We also found that the presence of ICE*Bs1* in cells caused a decrease in the formation of lysogens in a population exposed to SPß. We repeated the experiments described above with SPß::*spc* that is otherwise wild type and able to form lysogens. Lysogens were detected as spectinomycin-resistant colonies. When cells without ICE*Bs1* were infected with SPß::*spc (*MOI = 0.01), there was a 10-fold drop in both the optical density of the culture (Fig. 6A) and the number of CFUs (Fig. 6B). During the experiment, cells became lysogenized with SPß. Cells lysogenized by SPß are protected from killing by new SPß infection (Warner *et al.*, 1977). These lysogens continued to grow, and after about five hours the population of cells had increased and virtually all cells were SPß lysogens (Fig. 6B).

In contrast, when an ICE*Bs1*+ strain was infected with SPß::*spc* (MOI = 0.01), cell growth continued and there was no obvious drop in optical density (Fig. 6D) or the number of CFUs (Fig. 6E). Five hours after the initial infection, the number of phage in the culture was below the initial inoculum (Fig. 6E) and the number of SPß lysogens remained at approximately 10^4^ - 10^5^ per ml (Fig. 6F), a relatively small fraction of the total number of cells. Together, these results indicate that the presence of ICE*Bs1* in cells limits phage production, thereby protecting a population of cells from predation by SPß and limiting formation of SPß lysogens.

## Discussion

Results presented here demonstrate that ICE*Bs1* encodes an anti-phage system that inhibits production of the phage SPβ. This inhibition occurs upon *de novo* infection by SPβ and also upon induction of an SPβ lysogen. There is little or no direct effect on the formation of lysogens. The ICE*Bs1* gene *spbK* is both necessary and sufficient to inhibit SPβ: deleting *spbK* from ICE*Bs1* abolishes the phenotype, and expressing *spbK* in a strain missing ICE*Bs1* fully inhibits SPβ. Expression of *yonE* apparently triggers this anti-phage response, and co-expression of *spbK* and *yonE* in a strain that otherwise lacks ICE*Bs1* and SPβ rapidly kills the cells. We conclude that ICE*Bs1* encodes an abortive infection system that protects its host from predation by SPβ.

### Genes involved in protection against SPß

*yonE* is essential for the phage lytic cycle, but is not required for lysogen formation. Bioinformatic analysis of *yonE* and *yonF* revealed a possible role for these genes as components of the phage head-packing machinery, needed during the final stages of a phage’s lytic cycle.

SpbK contains a TIR domain. Where TIR domains have been studied they generally mediate protein-protein interactions through recognition of other TIR domains. However, examples of heterotypic interactions of TIR domains with non-TIR domain proteins have been described (Nimma *et al.*, 2017). In some bacterial pathogens, TIR-domain proteins modulate the host immune response (Newman *et al.*, 2006; Cirl *et al.*, 2008; Spear *et al.*, 2009; Alaidarous *et al.*, 2014). Recent work has implicated some bacterial TIR-domain proteins as being essential components of a class of anti-phage defense systems (“Thoeris”) (Doron *et al.*, 2018). Two of these Thoeris systems (from *Bacillus cereus* and *Bacillus amyloliquefaciens*) have been shown to non-specifically confer resistance to some myophages when reconstituted in *B. subtilis*, although the mechanism of this resistance is not understood.

Although SpbK and the Thoeris systems appear to have a common purpose, SpbK does not appear to be a component of a *B. subtilis* Thoeris antiphage system. The hallmark of Thoeris systems is a single gene encoding a NAD+ binding protein (ThsA) in proximity to (typically multiple) genes encoding TIR-domain proteins (ThsB) (Doron *et al.*, 2018). *spbK* is the only gene encoding a TIR-domain protein in *B. subtilis*. Furthermore, of all the genes in ICE*Bs1, spbK* is both necessary and sufficient for protection from SPβ, and there is no indication that SpbK contains a nucleotide binding domain. Irrespective of these differences, our analysis of SpbK raises the possibility that Thoeris anti-phage systems might function as abortive infection systems.

### Abortive infection systems

The anti-phage phenotype encoded by ICE*Bs1* resembles abortive infection systems that have been described for *Lactococcus*, *Escherichia coli*, and other bacteria. Such systems detect infection of a bacterial cell by phage and respond by inhibiting a host process needed for phage maturation and release (Labrie *et al.*, 2010). The mechanisms of inhibition vary widely, but often target a critical host process. The cellular process that is inhibited by SpbK is evidently essential for the host, as co-expression of *spbK* and *yonE* results in cell death. We have not yet determined what essential process (es) are targeted to cause SpbK-YonE-induced cell death.

### Cargo genes in ICEs

The cargo genes of mobile genetic elements, including ICEs, are those genes that are not necessary for the function of the mobile element, but are part of and transferred with the element. Cargo genes on a mobile element can often allow bacteria to rapidly acquire a new phenotype through acquisition of the element. Historically, most well studied ICEs were identified because of the phenotype (s) conferred by the cargo genes (Johnson and Grossman, 2015). Identification and characterization of the responsible genes revealed that they were in an ICE.

Many ICEs are being identified by bioinformatic analyses of sequenced bacterial genomes (Burrus *et al.*, 2002; Guglielmini *et al.*, 2011). In most of these analyses, it is not clear what, if any, phenotype is conferred by the ICE to its host. We suspect that many other ICEs with cargo genes of unknown function likely assist their hosts in mediating interactions with other mobile genetic elements.

## Methods

### Media and growth conditions

*E. coli* cells were grown in LB medium and on LB plates containing 1.5% agar at 37°C. *S. cereviseae* cells were grown in YPAD and on YPAD plates or synthetic dropout (SD) plates containing 2% agar and appropriate supplements to test for the indicated auxotrophies (James *et al.*, 1996; Gietz and Schiestl, 2007). *B. subtilis* cells were grown in LB or S7_50_ defined minimal medium with 0.1% glutamate (Jaacks *et al.*, 1989) with either glucose or arabinose (1% w/v) as a carbon source and on LB plates containing 1.5% bacto-agar or on Noble Agar (1.5%) for strains expressing both *spbK* and *yonE*. Antibiotics and other additives were used at the following concentrations for *E. coli*: carbenicillin (100 μg/ml), *B. subtilis*: kanamycin (5 μg/ml), spectinomycin (100 μg/ml), chloramphenicol (5 μg/ml). The Pspank(hy) promoter was activated with 1 mM isopropyl-ß-D-thiogalactopyranoside (IPTG), and the Pxyl promoter was activated with 1% (w/v) xylose, typically in cells grown in arabinose.

### Strains and alleles

#### SPß::*spc*

*spc* (spectinomycin resistance) was inserted between *yolB* and *yolC* in SPß. *spc* was amplified by PCR, and Gibson assembly (Gibson *et al.*, 2009) was used to join this fragment to genomic sequences containing the apparent bidirectional terminator located between the convergently transcribed genes *yolB* and *yolC* (de Hoon *et al.*, 2005) such that a copy of the terminator is located on each side of *spc*. This was then used to transform naturally competent *B. subtilis* cells selecting for resistance to spectinomycin. An antibiotic resistant strain (CMJ98) was identified and the location of the *spc* gene verified by sequencing. This strategy resulted in duplication of the terminator with *spc* located between the terminators. The resulting phage is referred to as SPß::*spc98*, or SPß::*spc*.

#### Δ(*spbK-rapI-phrI*)*1500*::*kan*

The region of ICE*Bs1* encoding *spbK*-*rapI*-*phrI* was replaced with *kan.* The deletion replaces all of *spbK*, *rapI*, and *phrI*, and was designed such that the orientation of *kan* and the *phrI* deletion boundary would be identical to the Δ(*rapI-phrI*)*342*::*kan* allele of IRN342 (Auchtung *et al.*, 2005).

#### Δ*yonEF396*::*spc*

A deletion-insertion that replaces *yonE* and *yonF* with a co-directional *spc* insertion.

#### Δ*yonE443*

The unmarked Δ*yonE443* allele was made by replacing *yonE* with the *cat* gene, flanked by *lox* sites (CMJ434). The Cre recombinase, expressed from the temperature-sensitive plasmid, pDR244, was then used to remove the *lox*-flanked *cat* gene by recombination. Strains were then cured of pDR244 by culturing them on LB + 1.5% agar at 45°C, as previously described (Meisner *et al.*, 2013; Johnson and Grossman, 2014), resulting in strain CMJ443.

#### Overexpression of *yonE*

The *yonE* coding sequence was cloned into a plasmid containing the IPTG-inducible Pspank(hy) promoter (Britton *et al.*, 2002), *lacI*, and either *spc* situated between genomic sequence from *amyE,* or *mls* situated between genomic sequence from *lacA*. The resulting construct was then transformed into competent *B. subtilis* cells. The following strains carrying a double-crossover of the given construct were identified by antibiotic resistance and PCR: CMJ403, *lacA*::{Pspank(hy)-*yonE lacI mls*}; CMJ616, *amyE*::{Pspank(hy)-*yonE lacI spc*}. Pspank(hy) is only partly repressed by LacI and was fully derepressed upon addition of 1 mM IPTG. Constructs lacking the *yonE* insert were also transformed into *B. subtilis* to generate the control alleles *lacA*::{Pspank(hy)-empty *lacI mls*} and *amyE*::{Pspank(hy)-empty *lacI spc*}.

#### Expression of *spbK*

To study expression of *spbK* in the absence of other ICE*Bs1* genes, a fragment containing the *spbK* coding sequence and 330 bp upstream was amplified by PCR and cloned into a plasmid for double-crossover integration into *lacA* or *amyE*. For cloning into *lacA*, the *spbK* fragment was cloned by Gibson assembly into a plasmid containing *kan* and parts of *lacA* suitable for double crossover. For cloning into *amyE*, the *spbK* fragment was cloned into a plasmid containing *cat* flanked by genomic sequences flanking the *amyE* locus by Gibson assembly. The resulting constructs were transformed into naturally competent *B. subtilis* cells and strains carrying a double crossover were identified as above, resulting in strains CMJ74 {*amyE*:: (*spbK cat*)} and CMJ684 {*lacA*:: (*spbK kan*)}.

### Plaque assays

To quantify the number of PFUs, samples with phage were diluted in LB and 100 μl of appropriate dilutions were mixed with 300 μl of an indicator strain at an OD600 of 0.5. Phage and cells were incubated at room temperature for 5 minutes, then mixed with 3 ml of soft agar (soft agar contains 10 g/l tryptone, 5 g/l yeast extract, 10 g/l NaCl, 6.5 g/l agar). The soft agar was spread on warm LB plates and incubated overnight at 37°C, allowing a lawn of cells to form. Plaques in the lawn were then counted. To photograph plaques, bacterial lawns were stained with 2,3,5 - triphenyltetrazolium chloride (Pattee, 1966) (TTC, Sigma). Briefly, 8 ml of 0.1% TTC in LB was pipetted onto plates and incubated at 37°C for 30 minutes. The TTC solution was then aspirated off and the petri dishes were photographed.

### Single-round infection experiments

Cells of the strain to be infected were cultured in rich medium to mid- to late exponential phase. The OD600 was then adjusted to 0.5 and 100 μl of cells was mixed with 10 μl medium containing 10^5^ PFU of SPß (MOI 1:100). Cells and phage were co-incubated for 5 min at 37°C, then washed 3 times by adding 1 ml LB, pelleting the cells in a microcentrifuge and removing the supernatant. The washed pellet was resuspended in 10 ml LB and incubated at 37°C with aeration to allow the phage to develop. Samples of the culture were taken at various time points and used immediately for plaque assays to quantify the concentration of infective centers (free phage + infected cells) in the culture.

### Quantification of lysogeny

The frequency of lysogenization was determined using SPß::*spc98*. 10^4^ PFUs of SPß::*spc98* in 10 μl LB were added to 100 μl of an indicator strain at an OD600 of 0.5 (an MOI of approximately 1:1000). Phage and cells were incubated at room temperature for 5 minutes, then cells were washed 3x with 1 ml LB to remove unbound phage. Cells were then spread on LB plates with spectinomycin to select for cells that had become lysongenized with SPß::*spc*.

### Yeast two-hybrid assays

The yeast two hybrid strains and vectors used in this study have been previously described (James *et al.*, 1996). Briefly, the coding sequence for *spbK* from amino acids 1-104 (N-terminus), 97-266 (TIR domain) and full length *spbK* were cloned into pGAD-C1 and fused to the *GAL4* activation domain or pGBDU-C3 and fused to the *GAL4* DNA binding domain. These vectors were then transformed into competent PJ69-4A cells using the LiAc method of Gietz and Schiestl (Gietz and Schiestl, 2007) and plated on synthetic dropout (SD) medium with appropriate supplements to select for acquisition of the plasmids. The ability to grow in the absence of leucine (pGAD-based plasmids) or uridine (pGBDU-based plasmids) was used to select clones that acquired each plasmid. To test for interaction between peptides, yeast strains carrying the plasmids of interest were spotted on SD medium and scored for growth in the absence of adenine, with growth indicating an interaction. As a control, strains carrying each individual plasmid were also scored for growth in the absence of adenine (all were negative).

### Mating assays

Mating assays were performed as previously described (Lee *et al.*, 2007; Johnson and Grossman, 2014). Briefly, donors and recipients were grown separately in minimal medium with 1% arabinose as a carbon source. RapI expression was induced in donors for 2 hours with 1% xylose. Approximately equal numbers of donors and recipients were then mixed, collected on a filter and placed on 1.5% agar plates buffered with Spizizens minimal salts (SMS agar contains 15 mM ammonium sulfate, 80 mM dibasic potassium phosphate, 44 mM monobasic potassium phosphate, 3.4 mM trisodium citrate, 0.8 mM magnesium sulfate, and 1.5% agar at pH 7.0) (Harwood and Cutting, 1990) for 90 minutes. Cells were rinsed off the filter, diluted, and spread on LB plates with selective antibiotics and incubated at 37°C overnight before quantification of colony forming units.

## Acknowledgements

We thank Mary Anderson and Josh Jones for helpful discussions and comments on the manuscript, T. DeWitt for preliminary experiments testing the effects of ICE*Bs1* on phages other than SPß, and Elena England for experiments confirming interactions between SPß and ICE*Bs1*.

Research reported here is based upon work supported, in part, by the National Institute of General Medical Sciences of the National Institutes of Health under award number R01GM050895 and R35 GM122538 to ADG. MMH was also supported, in part, by the NIGMS pre-doctoral training grant T32 GM007287. Any opinions, findings, and conclusions or recommendations expressed in this report are those of the authors and do not necessarily reflect the views of the National Institutes of Health.

---

**Table 2.** Yeast strains used.

| yeast strain* | genotype |
| --- | --- |
| CMJ620 | *S. cerevisiae* mat a ura3-52 leu2-3 his3 trp1 dph2Δ::HIS3 gal4Δ gal80Δ GAL2-ADE2 LYS2::GAL1-HIS3 met2::GAL-lacZ pCJ113 pCJ107 |
| CMJ621 | *S. cerevisiae* mat a ura3-52 leu2-3 his3 trp1 dph2Δ::HIS3 gal4Δ gal80Δ GAL2-ADE2 LYS2::GAL1-HIS3 met2::GAL-lacZ pCJ113 pCJ108 |
| CMJ622 | *S. cerevisiae* mat a ura3-52 leu2-3 his3 trp1 dph2Δ::HIS3 gal4Δ gal80Δ GAL2-ADE2 LYS2::GAL1-HIS3 met2::GAL-lacZ pCJ113 pCJ109 |
| CMJ626 | *S. cerevisiae* mat a ura3-52 leu2-3 his3 trp1 dph2Δ::HIS3 gal4Δ gal80Δ GAL2-ADE2 LYS2::GAL1-HIS3 met2::GAL-lacZ pCJ114 pCJ107 |
| CMJ627 | *S. cerevisiae* mat a ura3-52 leu2-3 his3 trp1 dph2Δ::HIS3 gal4Δ gal80Δ GAL2-ADE2 LYS2::GAL1-HIS3 met2::GAL-lacZ pCJ114 pCJ108 |
| CMJ628 | *S. cerevisiae* mat a ura3-52 leu2-3 his3 trp1 dph2Δ::HIS3 gal4Δ gal80Δ GAL2-ADE2 LYS2::GAL1-HIS3 met2::GAL-lacZ pCJ114 pCJ109 |
| CMJ632 | *S. cerevisiae* mat a ura3-52 leu2-3 his3 trp1 dph2Δ::HIS3 gal4Δ gal80Δ GAL2-ADE2 LYS2::GAL1-HIS3 met2::GAL-lacZ pCJ115 pCJ107 |
| CMJ633 | *S. cerevisiae* mat a ura3-52 leu2-3 his3 trp1 dph2Δ::HIS3 gal4Δ gal80Δ GAL2-ADE2 LYS2::GAL1-HIS3 met2::GAL-lacZ pCJ115 pCJ108 |
| CMJ634 | *S. cerevisiae* mat a ura3-52 leu2-3 his3 trp1 dph2Δ::HIS3 gal4Δ gal80Δ GAL2-ADE2 LYS2::GAL1-HIS3 met2::GAL-lacZ pCJ115 pCJ109 |

\* all yeast strains were derived from PJ69-4A (James *et al.*, 1996)

**Fig. S1.**
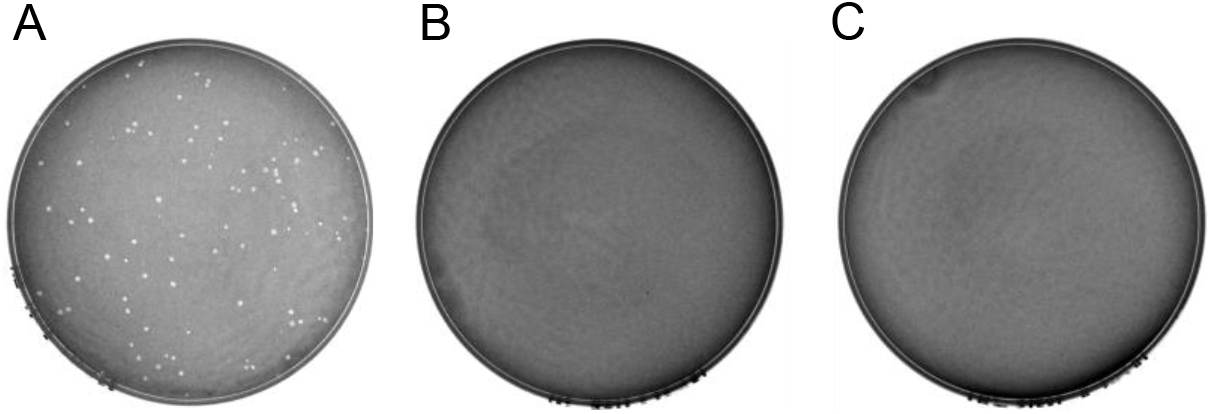
ICE*Bs1* prevents plaque formation by SPß. Various numbers of PFUs of SPß were mixed with the indicated strains, plated, incubated overnight and checked for the presence of plaques (methods). **A.** Approximately 100 PFUs of SPß were mixed with the indicator strain CU1050. **B.** Approximately 100 PFUs of SPß mixed with strain CMJ81 (the indicator strain CU1050 carrying ICE*Bs1*). **C.** Approximately 10^4^ PFU of SPß mixed with strain CMJ81 (the indicator strain CU1050 carrying ICE*Bs1*). In preliminary experiments, we found that ICE*Bs1* did not inhibit plaque formation by *B. subtilis* phages ø29, ø105, SP01, SP02, SPP1 or SP16. We conclude that the inhibitory effects of ICE*Bs1* are largely specific to SPß.

**Fig. S2.**
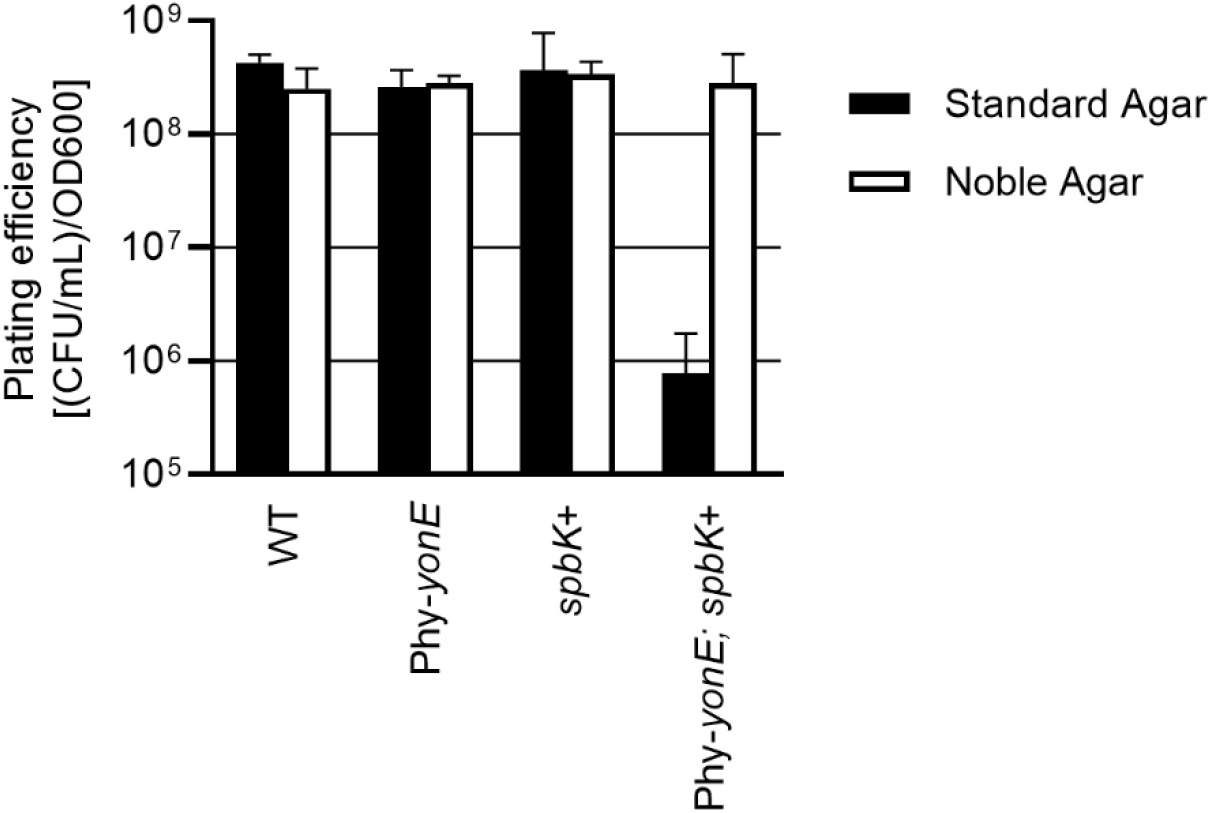
Co-expression of *spbK* and *yonE* results in a sensitivity to standard bacteriological agar. Strains null for ICE*Bs1* and SPß (PY79), expressing *yonE* (*amyE*::Phy-*yonE*, CMJ616), expressing *spbK* (*lacA*::*spbK*, CMJ684), or both *yonE* and *spbK* (CMJ685) were grown in minimal medium in the absence of IPTG. At an OD600 of 0.2, cultures were plated for CFUs on LB plates made with standard bacteriological agar (black bars) or on LB plates made with more rigorously purified Noble agar (white bars). Plating efficiency measured as CFUs/ml normalized to OD600.

**Fig. S3.**
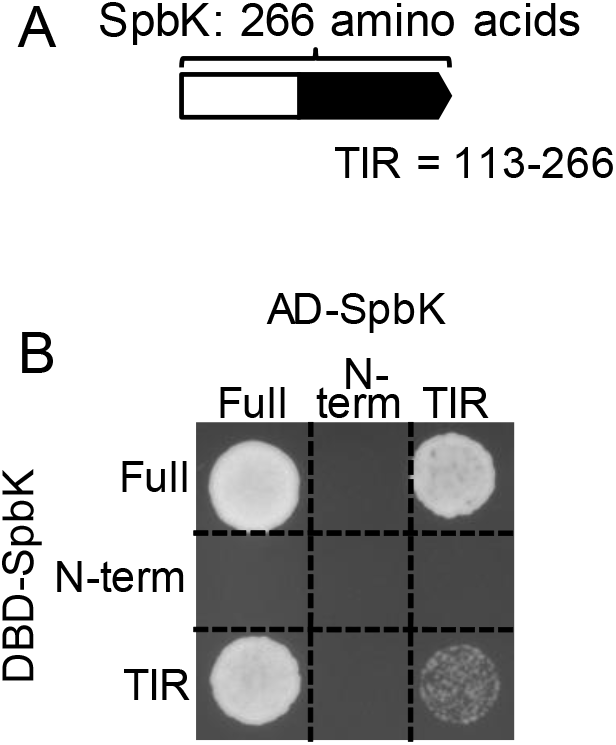
SpbK contains a TIR domain that mediates self-interaction. **A.** Map of the SpbK peptide sequence showing the location of the TIR domain in black. **B.** Yeast two-hybrid screen of SpbK fragments. Yeast strains carrying full length SpbK (Full), SpbK amino acids 1-104 (N-term) or SpbK amino acids 97-266 (TIR) bound to the *GAL4* DNA binding domain (DBD, Y-axis) and/or the *GAL4* activation domain (AD, X-axis) were spotted on medium selective for interaction between the bait and prey peptides and incubated at 30° C to allow for growth (methods). The following combinations were tested: AD-SpbK + DBD-SpbK (CMJ620), AD-N-term + DBD-SpbK (CMJ621), AD-TIR + DBD-SpbK (CMJ622), AD-SpbK + DBD-N-term (CMJ626), AD-N-term + DBD-N-term (CMJ627), AD-TIR + DBD-N-term (CMJ628), AD-SpbK + DBD-TIR (CMJ632), AD-N-term + DBD-TIR (CMJ633), AD-TIR + DBD-TIR (CMJ634).

## References

Abe, K., Takamatsu, T., and Sato, T. (2017) Mechanism of bacterial gene rearrangement: SprA-catalyzed precise DNA recombination and its directionality control by SprB ensure the gene rearrangement and stable expression of *spsM* during sporulation in *Bacillus subtilis*. Nucleic Acids Res 45: 6669–6683.

Alaidarous, M., Ve, T., Casey, L.W., Valkov, E., Ericsson, D.J., Ullah, M.O., et al. (2014) Mechanism of bacterial interference with TLR4 signaling by *Brucella* Toll/interleukin-1 receptor domain-containing protein TcpB. J Biol Chem 289: 654–668.

Auchtung, J.M., Aleksanyan, N., Bulku, A., and Berkmen, M.B. (2016) Biology of ICE*Bs1*, an integrative and conjugative element in *Bacillus subtilis*. Plasmid 86: 14–25.

Auchtung, J.M., Lee, C.A., Garrison, K.L., and Grossman, A.D. (2007) Identification and characterization of the immunity repressor (ImmR) that controls the mobile genetic element ICE*Bs1* of *Bacillus subtilis*. Mol Microbiol 64: 1515–1528.

Auchtung, J.M., Lee, C.A., Monson, R.E., Lehman, A.P., and Grossman, A.D. (2005) Regulation of a *Bacillus subtilis* mobile genetic element by intercellular signaling and the global DNA damage response. Proc Natl Acad Sci U S A 102: 12554–12559.

Barbe, V., Cruveiller, S., Kunst, F., Lenoble, P., Meurice, G., Sekowska, A., et al. (2009) From a consortium sequence to a unified sequence: the *Bacillus subtilis* 168 reference genome a decade later. Microbiology 155: 1758–1775.

Boratyn, G.M., Schäffer, A.A., Agarwala, R., Altschul, S.F., Lipman, D.J., and Madden, T.L. (2012) Domain enhanced lookup time accelerated BLAST. Biol Direct 7: 12.

Bose, B., Auchtung, J.M., Lee, C.A., and Grossman, A.D. (2008) A conserved anti-repressor controls horizontal gene transfer by proteolysis. Mol Microbiol 70: 570–582.

Britton, R.A., Eichenberger, P., Gonzalez-Pastor, J.E., Fawcett, P., Monson, R., Losick, R., and Grossman, A.D. (2002) Genome-wide analysis of the stationary-phase sigma factor (sigma-H) regulon of *Bacillus subtilis*. J Bacteriol 184: 4881–4890.

Burrus, V., Pavlovic, G., Decaris, B., and Guédon, G. (2002) The ICE*St1* element of *Streptococcus thermophilus* belongs to a large family of integrative and conjugative elements that exchange modules and change their specificity of integration. Plasmid 48: 77–97.

Carter, M.Q., Chen, J., and Lory, S. (2010) The *Pseudomonas aeruginosa* pathogenicity island PAPI-1 is transferred via a novel type IV pilus. J Bacteriol 192: 3249–3258.

Cirl, C., Wieser, A., Yadav, M., Duerr, S., Schubert, S., Fischer, H., et al. (2008) Subversion of Toll-like receptor signaling by a unique family of bacterial Toll/interleukin-1 receptor domain-containing proteins. Nat Med 14: 399–406.

Darmon, E., and Leach, D.R.F. (2014) Bacterial genome instability. Microbiol Mol Biol Rev MMBR 78: 1–39.

Delavat, F., Miyazaki, R., Carraro, N., Pradervand, N., and Meer, J.R. van der (2017) The hidden life of integrative and conjugative elements. FEMS Microbiol Rev 41: 512–537.

Doron, S., Melamed, S., Ofir, G., Leavitt, A., Lopatina, A., Keren, M., et al. (2018) Systematic discovery of antiphage defense systems in the microbial pangenome. Science 359.

Franke, A.E., and Clewell, D.B. (1981a) Evidence for conjugal transfer of a *Streptococcus faecalis* transposon (Tn*916*) from a chromosomal site in the absence of plasmid DNA. Cold Spring Harb Symp Quant Biol 45 Pt 1: 77–80.

Franke, A.E., and Clewell, D.B. (1981b) Evidence for a chromosome-borne resistance transposon (Tn*916*) in *Streptococcus faecalis* that is capable of “conjugal” transfer in the absence of a conjugative plasmid. J Bacteriol 145: 494–502.

Georgopoulos, C.P. (1969) Suppressor system in *Bacillus subtilis* 168. J Bacteriol 97: 1397–1402.

Gibson, D.G., Young, L., Chuang, R.-Y., Venter, J.C., Hutchison, C.A., and Smith, H.O. (2009) Enzymatic assembly of DNA molecules up to several hundred kilobases. Nat Methods 6: 343–345.

Gietz, R.D., and Schiestl, R.H. (2007) High-efficiency yeast transformation using the LiAc/SS carrier DNA/PEG method. Nat Protoc 2: 31–34.

Guglielmini, J., Quintais, L., Garcillán-Barcia, M.P., Cruz, F. de la, and Rocha, E.P.C. (2011) The repertoire of ICE in prokaryotes underscores the unity, diversity, and ubiquity of conjugation. PLoS Genet 7: e1002222.

Harwood, C.R., and Cutting, S.M. (1990) *Molecular biological methods for* Bacillus. Wiley, Chichester; New York.

Hochhut, B., Jahreis, K., Lengeler, J.W., and Schmid, K. (1997) CTnscr94, a conjugative transposon found in enterobacteria. J Bacteriol 179: 2097–2102.

Hoon, M.J.L. de, Makita, Y., Nakai, K., and Miyano, S. (2005) Prediction of transcriptional terminators in *Bacillus subtilis* and related species. PLoS Comput Biol 1: e25.

Jaacks, K.J., Healy, J., Losick, R., and Grossman, A.D. (1989) Identification and characterization of genes controlled by the sporulation-regulatory gene *spo0H* in *Bacillus subtilis*. J Bacteriol 171: 4121–4129.

James, P., Halladay, J., and Craig, E.A. (1996) Genomic libraries and a host strain designed for highly efficient two-hybrid selection in yeast. Genetics 144: 1425–1436.

Johnson, C.M., and Grossman, A.D. (2014) Identification of host genes that affect acquisition of an integrative and conjugative element in *Bacillus subtilis*. Mol Microbiol 93: 1284–1301.

Johnson, C.M., and Grossman, A.D. (2015) Integrative and conjugative elements (ICEs): what they do and how they work. Annu Rev Genet 49: 577–601.

Johnson, C.M., and Grossman, A.D. (2016) Complete genome sequence of *Bacillus subtilis* strain CU1050, which is sensitive to phage SPβ. Genome Announc 4.

Kunst, F., Ogasawara, N., Moszer, I., Albertini, A.M., Alloni, G., Azevedo, V., et al. (1997) The complete genome sequence of the gram-positive bacterium *Bacillus subtilis*. Nature 390: 249–256.

Labrie, S.J., Samson, J.E., and Moineau, S. (2010) Bacteriophage resistance mechanisms. Nat Rev Microbiol 8: 317–327.

Lazarevic, V., Düsterhöft, A., Soldo, B., Hilbert, H., Mauël, C., and Karamata, D. (1999) Nucleotide sequence of the *Bacillus subtilis* temperate bacteriophage SPbeta*c2*. Microbiol Read Engl 145 (Pt 5): 1055–1067.

Lee, C.A., Auchtung, J.M., Monson, R.E., and Grossman, A.D. (2007) Identification and characterization of *int* (integrase), *xis* (excisionase) and chromosomal attachment sites of the integrative and conjugative element ICE*Bs1* of *Bacillus subtilis*. Mol Microbiol 66: 1356–1369.

Lee, C.A., and Grossman, A.D. (2007) Identification of the origin of transfer (*oriT*) and DNA relaxase required for conjugation of the integrative and conjugative element ICE*Bs1* of *Bacillus subtilis*. J Bacteriol 189: 7254–7261.

Magot, M. (1983) Transfer of antibiotic resistances from *Clostridium innocuum* to *Clostridium perfringens* in the absence of detectable plasmid DNA. FEMS Microbiol Lett 18: 149–151.

Mays, T.D., Smith, C.J., Welch, R.A., Delfini, C., and Macrina, F.L. (1982) Novel antibiotic resistance transfer in *Bacteroides*. Antimicrob Agents Chemother 21: 110–118.

McHale, L., Tan, X., Koehl, P., and Michelmore, R.W. (2006) Plant NBS-LRR proteins: adaptable guards. Genome Biol 7: 212.

Meisner, J., Llopis, P.M., Sham, L.-T., Garner, E., Bernhardt, T.G., and Rudner, D.Z. (2013) FtsEX is required for CwlO peptidoglycan hydrolase activity during cell wall elongation in *Bacillus subtilis*. Mol Microbiol 89: 1069–1083.

Menard, K.L., and Grossman, A.D. (2013) Selective pressures to maintain attachment site specificity of integrative and conjugative elements. PLOS Genet 9: e1003623.

Narayanan, K.B., and Park, H.H. (2015) Toll/interleukin-1 receptor (TIR) domain-mediated cellular signaling pathways. Apoptosis Int J Program Cell Death 20: 196–209.

Newman, R.M., Salunkhe, P., Godzik, A., and Reed, J.C. (2006) Identification and characterization of a novel bacterial virulence factor that shares homology with mammalian Toll/interleukin-1 receptor family proteins. Infect Immun 74: 594–601.

Nimma, S., Ve, T., Williams, S.J., and Kobe, B. (2017) Towards the structure of the TIR-domain signalosome. Curr Opin Struct Biol 43: 122–130.

Nishi, A., Tominaga, K., and Furukawa, K. (2000) A 90-kilobase conjugative chromosomal element coding for biphenyl and salicylate catabolism in *Pseudomonas putida* KF715. J Bacteriol 182: 1949–1955.

Nugent, M.E. (1981) A conjugative “plasmid” lacking autonomous replication. J Gen Microbiol 126: 305–310.

Paik, S.H., Chakicherla, A., and Hansen, J.N. (1998) Identification and characterization of the structural and transporter genes for, and the chemical and biological properties of, sublancin 168, a novel lantibiotic produced by *Bacillus subtilis* 168. J Biol Chem 273: 23134–23142.

Pattee, P.A. (1966) Use of tetrazolium for improved resolution of bacteriophage plaques. J Bacteriol 92: 787–788.

Paulsen, I.T., Banerjei, L., Myers, G.S.A., Nelson, K.E., Seshadri, R., Read, T.D., et al. (2003) Role of mobile DNA in the evolution of vancomycin-resistant *Enterococcus faecalis*. Science 299: 2071–2074.

Perna, N.T., Plunkett, G., Burland, V., Mau, B., Glasner, J.D., Rose, D.J., et al. (2001) Genome sequence of enterohaemorrhagic *Escherichia coli* O157:H7. Nature 409: 529–533.

Ramsay, J.P., Sullivan, J.T., Stuart, G.S., Lamont, I.L., and Ronson, C.W. (2006) Excision and transfer of the *Mesorhizobium loti* R7A symbiosis island requires an integrase IntS, a novel recombination directionality factor RdfS, and a putative relaxase RlxS. Mol Microbiol 62: 723–734.

Rana, R.R., Zhang, M., Spear, A.M., Atkins, H.S., and Byrne, B. (2013) Bacterial TIR-containing proteins and host innate immune system evasion. Med Microbiol Immunol (Berl) 202: 1–10.

Rao, V.B., and Feiss, M. (2015) Mechanisms of DNA packaging by large double-stranded DNA viruses. Annu Rev Virol 2: 351–378.

Rauch, P.J., and De Vos, W.M. (1992) Characterization of the novel nisin-sucrose conjugative transposon Tn*5276* and its insertion in *Lactococcus lactis*. J Bacteriol 174: 1280–1287.

Ravatn, R., Studer, S., Springael, D., Zehnder, A.J.B., and Meer, J.R. van der (1998) Chromosomal integration, tandem amplification, and deamplification in *Pseudomonas putida* F1 of a 105-kilobase genetic element containing the chlorocatechol degradative genes from *Pseudomonas* sp. strain B13. J Bacteriol 180: 4360–4369.

Roberts, M.C., and Smith, A.L. (1980) Molecular characterization of “plasmid-free” antibiotic-resistant *Haemophilus influenzae*. J Bacteriol 144: 476–479.

Sakaguchi, Y., Hayashi, T., Kurokawa, K., Nakayama, K., Oshima, K., Fujinaga, Y., et al. (2005) The genome sequence of *Clostridium botulinum* type C neurotoxin-converting phage and the molecular mechanisms of unstable lysogeny. Proc Natl Acad Sci U S A 102: 17472–17477.

Sebaihia, M., Wren, B.W., Mullany, P., Fairweather, N.F., Minton, N., Stabler, R., et al. (2006) The multidrug-resistant human pathogen *Clostridium difficile* has a highly mobile, mosaic genome. Nat Genet 38: 779–786.

Shoemaker, N.B., Smith, M.D., and Guild, W.R. (1980) DNase-resistant transfer of chromosomal *cat* and *tet* insertions by filter mating in *Pneumococcus*. Plasmid 3: 80–87.

Smith, J.L., Goldberg, J.M., and Grossman, A.D. (2014) Complete genome sequences of *Bacillus subtilis* subsp. subtilis laboratory strains JH642 (AG174) and AG1839. Genome Announc 2 https://www.ncbi.nlm.nih.gov/pmc/articles/PMC4082004/. Accessed January 6, 2020.

Spear, A.M., Loman, N.J., Atkins, H.S., and Pallen, M.J. (2009) Microbial TIR domains: not necessarily agents of subversion? Trends Microbiol 17: 393–398.

Stuy, J.H. (1980) Chromosomally integrated conjugative plasmids are common in antibiotic-resistant *Haemophilus influenzae*. J Bacteriol 142: 925–930.

Warner, F.D., Kitos, G.A., Romano, M.P., and Hemphill, H.E. (1977) Characterization of SPβ: a temperate bacteriophage from *Bacillus subtilis* 168M. Can J Microbiol 23: 45–51.

Wozniak, R.A.F., and Waldor, M.K. (2010) Integrative and conjugative elements: mosaic mobile genetic elements enabling dynamic lateral gene flow. Nat Rev Microbiol 8: 552–563.

Youngman, P., Perkins, J.B., and Losick, R. (1984) Construction of a cloning site near one end of Tn*917* into which foreign DNA may be inserted without affecting transposition in *Bacillus subtilis* or expression of the transposon-borne *erm* gene. Plasmid 12: 1–9.

Zahler, S.A., Korman, R.Z., Rosenthal, R., and Hemphill, H.E. (1977) *Bacillus subtilis* bacteriophage SPbeta: localization of the prophage attachment site, and specialized transduction. J Bacteriol 129: 556–558.

Zyl, L.J. van, Sunda, F., Taylor, M.P., Cowan, D.A., and Trindade, M.I. (2015) Identification and characterization of a novel *Geobacillus thermoglucosidasius* bacteriophage, GVE3. Arch Virol 160: 2269–2282.

